# Molecular Characterization of the Progressive Landscape of Depression

**DOI:** 10.64898/2026.05.22.727217

**Authors:** Vandana Sharma, Elena Payne, Samuel Gonzalez Garcia, Lauren Fang, Kiran Boyinepally, Akiko Sumitomo, Toshifumi Tomoda, David A. Lewis, Robert Mccullumsmith, Etienne Sibille, Rammohan Shukla

## Abstract

Major Depressive Disorder (MDD) frequently follows a recurrent trajectory of episodes and remissions, often culminating in treatment-resistance. Molecular differences defining state-specific changes during episode and remission have been explored. However, progressive differences—defined here as cross-sectional linear trends across clinical stages from first to recurrent episodes or remissions, reflecting increasing illness burden over time—remain poorly understood, limiting sustained therapeutic outcomes. Here, we analyzed RNA-seq data from postmortem sgACC to identify progressive differences across MDD episodes or remission relative to state-specific differences, using an integrative assessment of molecular and cellular specificity, genetic-risk, disease-comorbidity and potential therapeutic targets. Differential expression analysis showed greater overlap between progressive and state-specific differences during remission than episode. Pathway enrichment highlighted disruptions in extracellular-matrix pathways shared by state-specific and progressive episodes, while metabolic and catalytic pathways were restored during remission. Cell-type-specific analyses showed that progressive changes were linked to superficial-layer intra-telencephalic neurons, whereas state-specific changes were enriched in pyramidal neuron subtypes and deeper layer SST-positive interneurons. Genome-wide association-informed enrichment analysis further linked these transcriptomic changes to genetic risk factors and symptom dimensions. Anhedonia was associated with both state-specific episode and progressive-remission signatures, suggesting that it is a persistent trait-like feature of MDD. Finally, an integrative pharmacological analysis revealed shared molecular mechanisms between pro-disease and therapeutic targets, highlighting pleiotropic effects of key pathways depending on disease state and dosage. Together, these findings provide a novel perspective on biological underpinnings of MDD progression over episodes or remissions and identify pharmacological targets that account for pathological and/or compensatory/therapeutic processes.

## Introduction

Major Depressive Disorder (MDD) cost the US economy over $326.2 billion in 2023, including direct costs, such as healthcare expenditures and medications and indirect costs, such as lost productivity, disability, and absenteeism [1]. Clinical evidence indicates that MDD typically follows a periodic trajectory of recurring depressive episodes, characterized by increasing severity, longer duration, and increasing resistance to antidepressant therapy [2, 3]. These episodes are interspersed with progressively shorter and incomplete remission phases often leading to treatment-resistance depression and reduced functional capacity. Addressing MDD requires a comprehensive understanding of the molecular and cellular factors, their genetic associations, and the complex interactions among them that drive disease process and influence its trajectory.

In our previous report [4], we analyzed transcriptomic data from the postmortem subgenual anterior cingulate cortex (sgACC), a key integration center of mood and reward pathways critically implicated in MDD [5–7]. Our analysis was based on the premise that the clinical states (episode or remission) of MDD is associated with distinct “state-dependent” molecular changes. We examined individuals who died during distinct phases of the MDD trajectory (Figure 1A), including the first depressive episode, first remission, recurrent episode, and recurrent remission. This analysis revealed clear patterns of molecular changes associated with disease states with sets of genes whose expression oscillated between episode and remission phases. These changes were mostly related to synaptic functions, bioenergetics, inflammation and immune activation [4]. While these comparisons effectively highlighted the dynamic differences between episode and remission states, it remains unclear whether a distinct set of gene expression changes accumulate over time, contributing to disease progression and worsening across episodes. This dataset presents an opportunity to address this by examining stage-wise (cross-sectional) trends, modeled as progressive changes from control to first episode/remission and from there to recurrent episodes/remissions.

**Figure 1.**
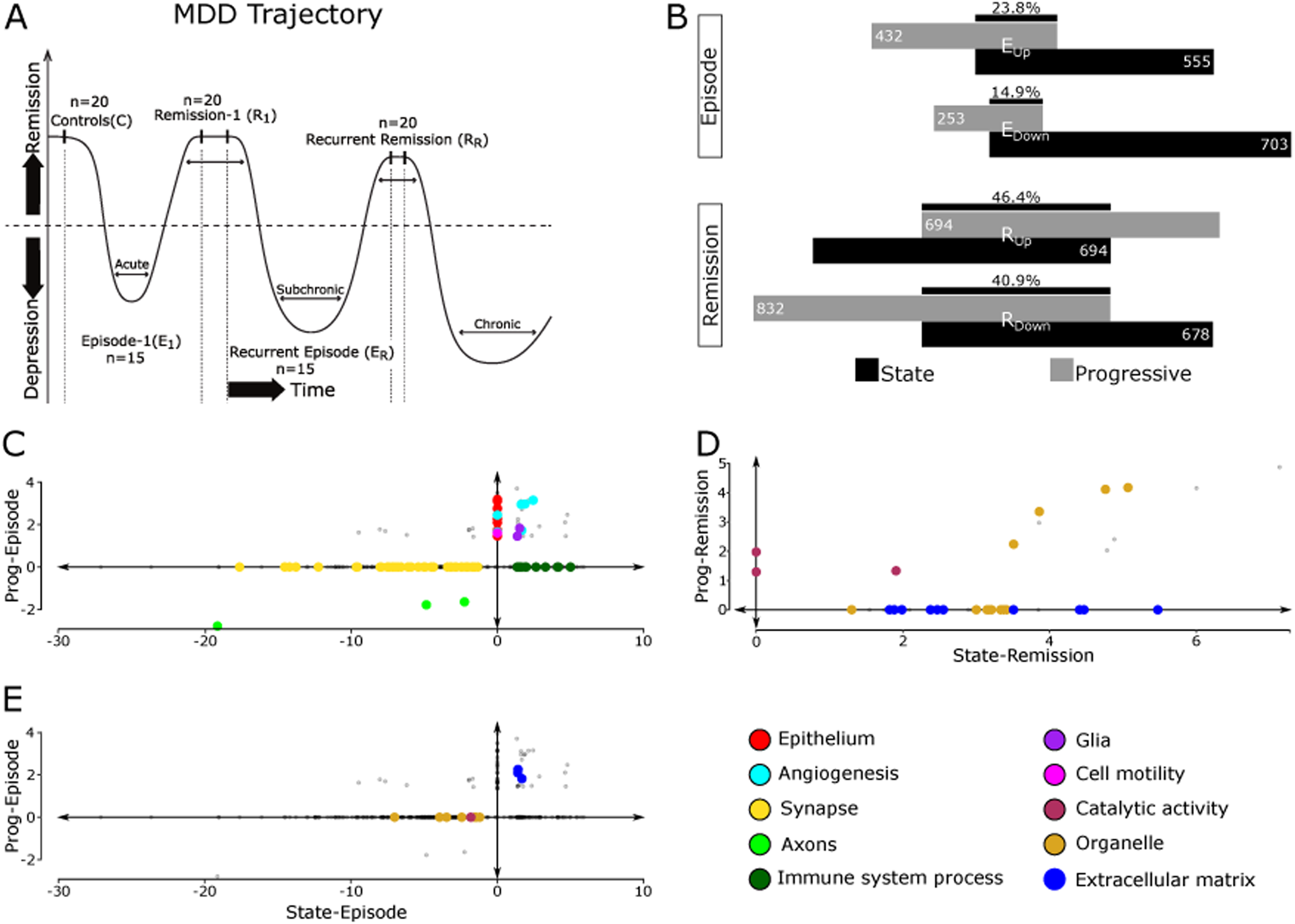
Transcriptomic and pathway alterations in sgACC across MDD phases: **(A)** Study design and sample groups overlaid on a schematic of MDD clinical progression, depicting distinct disease phases. **(B)** Similarity of DEGs for episode [E] and remission [R] phases between state and progressive contrasts. State contrasts compare controls with all episode or remission cases to capture disease state–dependent changes. Progressive contrasts model cross-sectional linear trends across clinical stages (Control → E1 → ER; Control → R1 → RR) to identify genes varying consistently with illness stage. Black bars indicate the Jaccard similarity index (intersection/union x 100) for each comparison. Overlap counts (intersection/union): E_Up_ = 189/796, E_Down_ = 184/832, R_Up_ = 440/948, R_Down_ = 439/1061. **(C–E)** Mapping of pathway profiles (q-value < 0.01) onto Cartesian coordinates, with each axis representing the significance score calculated as -log10(q-value) and pathways classified into functional themes (color-coded). (E) plots the episode contrast of the significant remission pathways shown in (D). Note that ECM-related pathways upregulated during remission (D), are also upregulated during episodes (E). See Supplementary Table S3 for DEGs associated with each contrast and Supplementary Table S4 for all pathways associated with each contrast.

We hypothesized that the accumulation of subthreshold molecular changes during disease progression, combined with residual effects from depressive symptoms persisting during remission and suggesting incomplete remission, contributes to MDD chronicity and exacerbation over time and episodes. We predicted a unique set of genes and biological pathways would be associated with disease progression, independent of state-dependent changes, suggesting novel targets for therapeutic interventions aimed at the disease trajectory.

## Materials and methods

### Human brain samples

Postmortem sgACC tissue was obtained as previously described [4]. Briefly, brain samples were collected during routine autopsies conducted at the Allegheny County Medical Examiner’s Office (Pittsburgh PA), following protocols approved by the University of Pittsburgh’s Committee for Oversight of Research and Clinical Training Involving Decedents and Institutional Review Board for Biomedical Research. DSM-IV diagnoses were determined by an independent panel of clinicians based on structured interviews with family members, clinical records, toxicology reports, and neuropathology exams. Control subjects were similarly evaluated to confirm the absence of active psychiatric or neurological disorders at the time of death. A small number of control cases had previous DSM-IV diagnosis (adjustment disorder with depressed mood, past) and were on non-psychiatric medications at death.

However, neuropathological assessment revealed no evidence of brain disease, and these cases were retained as controls. The study included 90 samples: 20 control individuals, 20 individuals experiencing a first MDD episode (E1), 15 in remission after a single episode (R1), 20 with recurrent episodes (ER), and 15 in remission following multiple episodes (RR) (Supplementary Table 1 & 2). Wherever possible, efforts were made to minimize differences in age, postmortem interval (PMI), brain pH, and RNA integrity number (RIN) across cohorts. Information on cause and mode of death, psychotropic medication use, substance dependence, and history of psychosis was recorded. Coronal sections encompassing all six cortical layers were collected as previously described [8]. Samples from this cohort have been utilized in earlier large-scale transcriptomics [4] and proteomics [9] studies.

### RNA isolation and library construction

Total RNA was isolated from homogenized samples using the RNeasy mini kit (Qiagen, Cat. No. 74104), incorporating an in-column DNase treatment with RNase-Free DNase (Qiagen, Cat. No. 79254) to eliminate genomic DNA contamination. Sequencing libraries were generated using the SMARTer stranded total RNA-seq kit (Clontech Laboratories, Cat. No. 634876). All procedures were carried out following the manufacturer’s guidelines.

### Sequencing and data processing

Sequencing libraries were processed on the Illumina HiSeq2500 platform, generating 2 × 100 bp paired end reads. On average, 41 million paired-end reads were obtained per sample. As described in our previous work [10] these reads were aligned to the human reference genome (GRCh38) from Ensembl using the HISAT2 aligner.

Read count data were subsequently generated for exons and transcripts using the *GenomicFeatures* and *GenomicAlignments* packages in R, along with the gene model (GTF file) provided by Ensembl. On average, 9.7 million uniquely mapped reads per sample aligned to approximately 17,000 genes, which were used for downstream analyses.

### Differential expression analysis

We focused on identifying progressive transcriptomic changes, defined as cross-sectional linear trends across clinical stages from control to first and then recurrent episodes or remissions (see Supplementary Figure S1, notes). Both analyses were performed using DESeq2 package in R [11], which is well suited for datasets with modest or uneven replicate numbers across groups, as in the present study. Control, single-episode, and recurrent-episode (or remission) groups were treated as ordinal variables (1, 2, 3) to model group-level linear trends across clinical stages. State contrasts were analyzed as previously described [4], comparing controls to all individuals in MDD episode or to all individuals in remission. However, as detailed in the section below, the present analysis additionally incorporated several covariates into the DESeq2 model to better control for potential confounding effects. After removing low-expressed genes (mean row-sum ≤ 5), 17,175 and 16,456 genes were included in the progressive and state analysis, respectively (Supplementary Table 3).

To assess the variance in gene expression associated with clinical and demographic variables, we performed a variance partitioning analysis across all samples in the cohort using the variancePartition package in R [12]. Batch, RNA ratio, PMI, brain pH, age, and RIN were among the top-ranked variables, while medications at the time of death, mode of death, race, and sex were among the bottom-ranked variables in terms of average variance explained (Supplementary Figure S2A). Although these variables accounted for lower average variance, they still influenced the expression of several genes with potential biological relevance. Previous studies have included medication at the time of death as a covariate in transcriptomic analyses due to its potential impact on gene expression [13]. Therefore, we regressed out all these variables by incorporating them into the design matrix in DESeq2 as covariates: design = ∼Batch + RNA ratio + PMI + Brain pH + Age + RIN + Medication + Mode of Death + Race + Sex + Condition. Here, Condition represents the primary variable of interest, corresponding to either progressive or state changes. To ensure that covariate correction did not over-adjust or remove biologically meaningful variance, we assessed collinearity between covariates and found that no variables were highly correlated (Supplementary Figure S2B). While no gold standard exists for validating the effectiveness of covariate correction in this context, we applied two indirect approaches. First, we examined the p-value histogram, a widely used diagnostic tool for evaluating the distribution of statistical significance [14]. Second, we examined pathway profiles and disease-based enrichment associated with the differentially expressed genes. We used the fdrtool package in R to model the distribution of raw p-values and estimate local false discovery rate. The histograms (Supplementary Figure S2C) display raw p-values following fdrtool adjustment, which showed a uniform distribution, consistent with well-calibrated p-values. As discussed in the results section, several MDD-relevant pathways emerged in the analysis. A p-value threshold of 0.05 was used.

### Pathway enrichment analysis

Biological pathways affected in both progressive and state contrasts were identified using Gene Ontology (GO) analysis [15] via the GO web interface [https://geneontology.org/]. Differentially expressed genes (DEGs) for each contrast were tested for their overrepresentation in the three GO categories—Biological Process (GOBP), Molecular Function (GOMF), and Cellular Component (GOCC)—using Fisher’s Exact test, with significant pathways identified at q-value < 0.05. Separate analyses were performed for up-and downregulated genes in each contrast. To facilitate comparison and visualization while maintaining directional identity, q-values were transformed into significance scores (SS) using - log_10_(q-value), with downregulated SS multiplied by -1 to distinguish them from upregulated scores. The final values were merged into a single contrast output and plotted.

To summarize the ontologies and better characterize the biological changes in the enrichment results, we performed a focused analysis around predefined functional theme (color-coded in Figure 1). Specifically, selected ontologies representing themes of interest were manually chosen (referred to as parent terms), and their associated child terms were identified from the significant GO terms linked to both contrasts using the GO.db package in R. The package utilizes specific functions to retrieve all child terms linked to higher-order GO terms (parent-term) based on their hierarchical structure.

### Quantitative polymerase chain reaction (qPCR

To ensure technical validation and concordance with RNA-seq findings, we confirmed expression differences using a qPCR-based assay on the same RNA samples. Genes for validation (*ARID5B* and *DDX25*) were selected based on their RNA-seq differential expression profiles and their presence within the subset of differentially expressed genes that overlapped with and drove the enriched GO terms in the progressive episode and progressive remission contrasts. To maximize sensitivity for detecting concordance between the two techniques, the top six samples representing either a control or diseased state were selected based on those exhibiting the most pronounced and consistent separation in normalized RNA-seq expression values of the selected genes. Genes associated with the state contrast were previously validated in our earlier study using this cohort [4].

Total RNA, identical to that used for sequencing library preparation (concentration range: 8.4-10.8 ng/μl), was reverse transcribed into cDNA using PrimeScript RT master mix (TaKaRa), with 50 ng of total RNA in a final reaction volume of 10μl. The reaction mixture, containing cDNA, primers, and TB Green Premix Ex Taq (Tli RNaseH Plus) (TaKaRa), was prepared in a 96-well PCR plate, and qPCR was performed in triplicate using the CFX96 Real-Time System (Bio-Rad) using a two-step cycling protocol (95°C for 15 sec followed by 60°C for 30 sec), repeated for 40 cycles. Results were normalized to *GAPDH* as an internal control. To assess significance, we fitted a linear regression model on normalized expression values across cohorts representing the progressive episode and remission groups, consistent with the treatment of ordinal group variables in the progressive contrast of the differential expression analysis. Primers used are detailed below

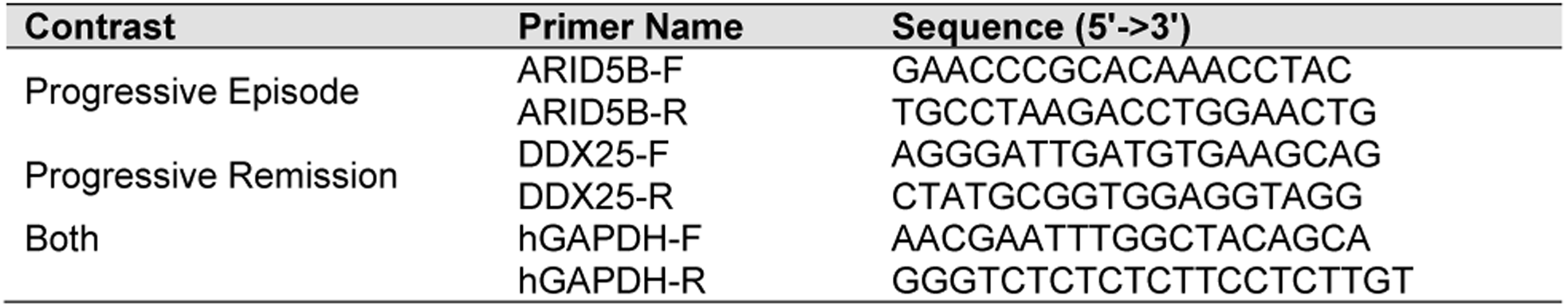

### Hypergeometric analysis

To assess the association of the progressive and state contrasts with sgACC cell types, diseases and drugs, we performed hypergeometric overlap analysis using the GeneOverlap package in R [16]. This approach tests the significance of overlap between gene sets using a hypergeometric distribution, with a genomic background of 21,196 genes (default in the GeneOverlap package). Similar to GO term overrepresentation analysis, GeneOverlap treats this as a statistical test of independence between gene sets, applying Fisher’s exact test to determine significance. Unless otherwise specified, results are reported at q-value < 0.05.

Drugs and disease-associated gene sets were obtained from Enrichr gene set libraries in .gmt format. The drug-specific gene sets originate from various transcriptomic studies on mice and human cell-lines treated with different drugs with known mechanisms of action (also see notes for supplementary table S8). The disease-specific gene sets were sourced from DisGeNET—a curated database integrating gene-disease associations from multiple sources, including the literature and expert-reviewed repositories [17]. Additionally, we expanded the disease-specific gene sets using the PsyGeNET database, which focuses on gene sets specifically linked to psychiatric disorders [18].

### Cell-type proportion estimation and enrichment analysis

To estimate cell-type proportions and identify putative disease-associated cellular differences, we performed deconvolution analysis. The analysis utilized two input datasets: bulk RNA-sequencing expression data from the present study, representing a mixture of cell types across both state and progressive contrasts, and single-cell expression data from the Allen Brain Atlas [19], which served as a reference for estimating cell-type proportions in the bulk data. Cell-type proportion estimation was conducted using a support vector regression approach, as described by Newman et al [20], and implemented in the granulator package in R.

To assess the significance of altered cell-type proportions during progressive changes, we fitted a linear regression model for each cell type, using cell-type proportion as the dependent variable, and ordered conditions (control, episode 1, and recurrent episodes or control, remission 1, and recurrent remission) as the predictor, similar to the treatment of ordinal variable in the progressive contrast of both the differential expression and qPCR analysis. The trend p-value was extracted from each model to determine whether a cell type proportion exhibited a significant linear shift across conditions. Likewise, groupwise comparisons between control and all subjects in either episodes or remissions were conducted following the state contrast framework in the differential expression analysis, and a two-sample t-test was performed for each cell type to evaluate differences in proportions between the episode and remission state groups.

To validate our approach, we repeated all steps using an ordinary least square–based deconvolution method [21] implemented in the granulator package. Since no significant changes in cell-type proportions were observed across contrasts using either method, we further examined the enrichment of sgACC cell-type–specific marker genes within the DEGs of all contrasts using a hypergeometric overlap test, as described above. The cell-type–specific gene sets were obtained from recent sgACC-specific single-cell analyses [22]. All enrichments were reported at q-value < 0.05.

### MAGMA Gene-set analysis

To assess the genetic component associated with state and progressive contrasts, we performed gene-set enrichment analysis using MAGMA (Multi-marker analysis of GenoMic Annotation; version: v1.10) [23] on meta-analysis summary statistics from two recent MDD genome-wide association studies, publicly available through the Psychiatric Genomics Consortium [24]. The first study included individuals from 29 countries across diverse and admixed ancestries [25], while the second study integrated MDD ascertainment criteria and symptom structures [26].

Gene-set enrichment analysis was conducted separately for up- and downregulated DEGs associated with state and progressive contrasts, using both competitive and self-contained testing approaches. The competitive analysis tested whether each DEG set was more strongly associated with MDD than other genes across the genome, while the self-contained analysis assessed whether each DEG set showed a significant association with MDD independent of other genes. Adjustments were made for gene size, gene set density, and linkage disequilibrium (LD) structure. Results are reported as both nominal and FDR-corrected p-values.

## Data availability

All datasets analyzed during the current study are available as supplementary tables. Raw data (count matrix, fastq.gz, or .bam) are available from the corresponding author on reasonable request.

## Results

### Distinct gene expressions and pathway changes are associated with MDD progression over episodes or remission compared to disease states

RNAseq-based expression profiles were analyzed in postmortem sgACC samples, comprising one control group (C) and four cohorts representing different phases of MDD: first depressive episode (E_1_), remission after the first episode (R_1_), recurrent episode (E_R_), and remission after recurrent episodes (R_R_) (Figure 1A). We performed differential expression analyses to examine two main types of contrasts representing disease states (episode or remission) and progressive changes across episodes or remission phases (See methods, Supplementary Figure S1), using a nominal significance threshold of p-value < 0.05 across all contrasts. The “state” contrasts were partially explored in our previous work, where control samples were compared with combined episode (E_1_ and E_R_) or remission (R_1_ and R_R_) groups (referred to as state-episode and state-remission, respectively). Here, we expand on this framework by integrating progressive contrasts—fitting a linear model to identify genes with progressive up- and down-regulation across cohorts C, E_1_, and E_R_, (referred to as progressive episode) and across cohorts C, R_1_, and R_R_ (referred to as progressive remission).

Episode- and remission-specific DEGs showed minimal overlap between progressive and state contrasts, highlighting transcriptional distinctness (Supplementary Figure S3). Episode-specific DEGs showed moderate overlap between progressive and state contrasts (Figure 1B; 23.8% upregulated, 14.9% downregulated), while remission-specific DEGs showed substantially higher overlap (Figure 1B; 46.4% upregulated, 40.9% downregulated). These results suggest that episode-related signatures may differ more between current disease burden and illness progression, whereas remission-associated signatures appear relatively stable across contrasts. Together these patterns point to potentially distinct molecular mechanisms underlying the depressive state and its progression during episodes.

To better separate subtle progressive changes in the context of more pronounced state changes, we mapped the significance scores of the GO-terms with q-value < 0.01 onto Cartesian coordinates and classified them into different themes (Figure 1, color codes). The Y-axis represents progressive episodes (Figure 1C & E) or remission (Figure 1D), while the X-axis represents state-episodes (Figure 1C & E) or state-remission (Figure 1D). Dots along the axes represent biological changes unique to one contrast, whereas off-axis dots indicate shared changes captured by both contrasts.

For the episode phases (Figure 1C), several changes were exclusive to either the progressive or state contrasts. For example, pathways associated with epithelial cell activity (red) and cell-motility (pink) were upregulated during the progressive episode. In contrast, and consistent with prior findings, downregulation of synapse-related pathways (yellow) and upregulation of immune-related pathways (dark green) were specific to the state contrasts and did not exhibit progressive changes across episodes. Other pathways were shared across contrast. Upregulated pathways related to angiogenesis (cyan) and glial activity (purple) were more closely linked to the progressive contrast but also present in state contrast. In addition, downregulated pathways linked to axonal function (light green) were shared between the progressive and state contrasts, suggesting that disrupted sgACC output due to axonal downregulation accumulates over the course of MDD.

For the remission phases (Figure 1D), only a few upregulated pathways were common to both progressive and state contrasts. For instance, pathways associated with catalytic (maroon) and organelle activity (tan) were upregulated in both contrasts. However, pathways associated with extracellular matrix (ECM, dark blue) were exclusively upregulated during state-remission. Mapping these remission-associated pathways to progressive or state contrasts for episode (Figure 1E) revealed that changes in catalytic and organelle activity pathways were absent during progressive episodes but downregulated during the state-episode, suggesting a restoration of cellular and metabolic pathways during remission. Interestingly, ECM-related pathways, were also upregulated in both the progressive and state contrasts for episode, indicating irreversible alterations associated with this pathway throughout the MDD trajectory.

As a technical validation, qPCR showed strong concordance (Supplementary Figure S4) for two DEGs—*ARID5B* and *DDX25*—associated with progressive episodes and remission-related pathways, respectively.

Taken together, these findings highlight distinct gene expression and pathways changes associated with MDD progression over episodes or remission compared to disease states. Some biological changes cycle with episodes without progressing overtime, particularly those involving synapse and signaling. In contrast, alterations in ECM pathways and restoration of catalytic and organelle-related functions progress over episodes and appear irreversible during remission.

### MDD states and progression target specific cell-types without altering cell proportions

To investigate putative cell-type specific changes, we utilized available single nucleus RNA-seq data from the ACC [19] and performed deconvolution analysis to estimate changes in cell-type proportion (see methods). No significant changes were observed for either progressive or state contrasts during episodes and remission (supplementary table 5 and 6), suggesting stable neuronal composition throughout the disease.

Next, since MDD may alter cell function rather than their generation or death, we performed DEG cell-specific enrichment analysis (q-value < 0.05). Among the downregulated DEGs (Figure 2, bottom), only episodic phase linked to both state and progressive contrast showed cell-type specific enrichment. State-episode exhibited the most prominent DEG cell-type specific enrichment, predominantly affecting excitatory pyramidal neurons, including intra-telencephalic (IT) and extra-telencephalic (ET) neurons across all layers, as well as near-projecting (NP) neurons in deeper layers. Among interneurons, consistent with our previous studies [27, 28], state-episode DEG changes were specifically linked to SST-positive neurons in layers 5-6. Progressive DEG changes during episode on the other hand were linked exclusively to IT pyramidal neurons of layer 3, suggesting that cortico-cortical connections – the function of IT neurons [29, 30], are progressively disrupted during disease progression. Among the upregulated DEGs (Figure 2, top), only state contrasts linked to both episode and remission phase showed exclusive enrichment of DEGs in endothelial cells, which are regulators of the ECM—a pathway dysregulated in the functional pathway analysis described above (Figure 1D & E).

**Figure 2.**
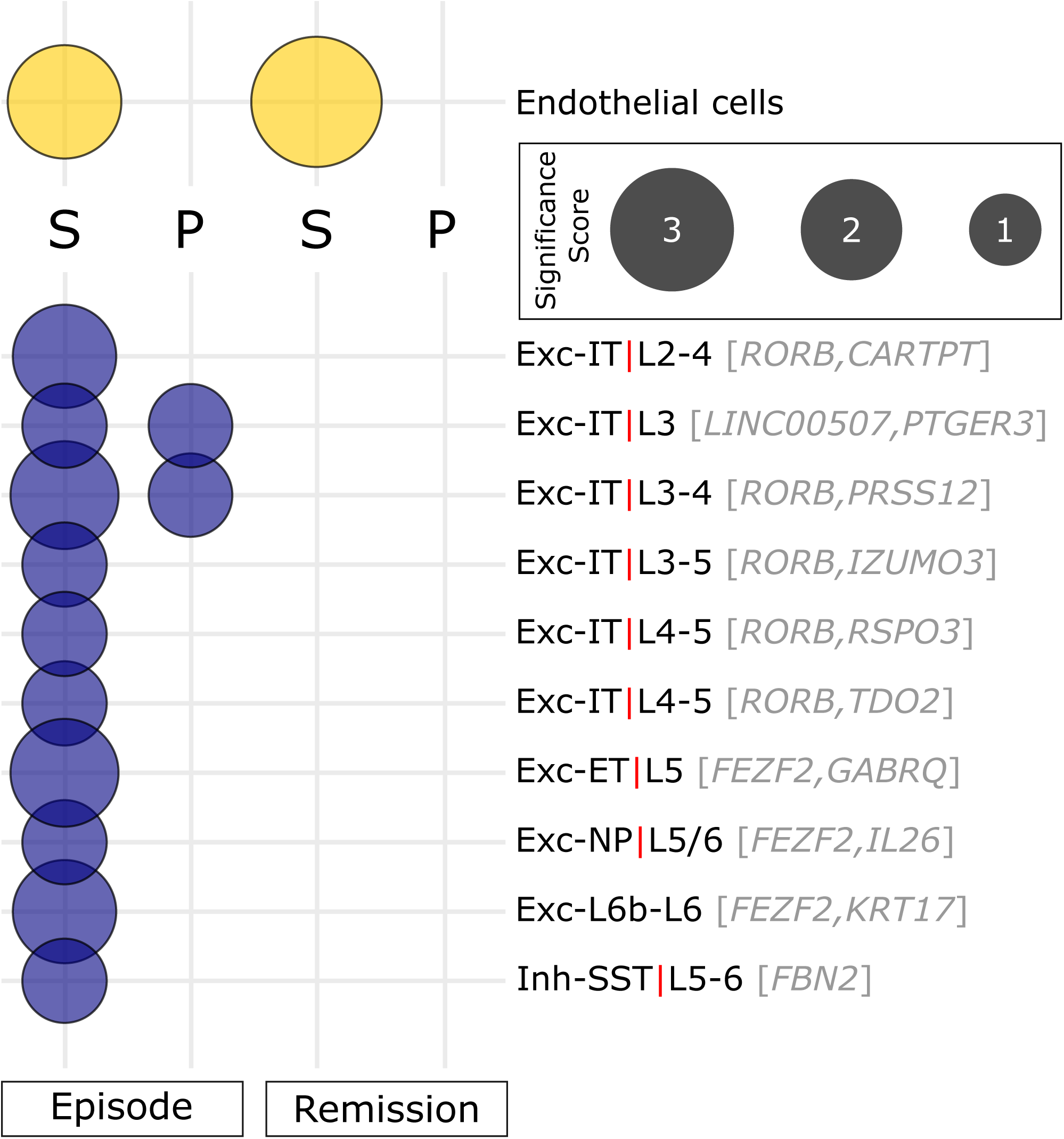
Cell-type-specific association across MDD phases: Cell-type-specific enrichment of downregulated (blue) and upregulated (yellow) genes. The circle size is proportional to the significance score, calculated as -log_10_(q-value). For excitatory (Exc) and inhibitory (Inh) neurons, their classification (IT, ET, NP, and SST), layer-specific distribution, and top marker genes are shown.

Together, the results do not support cell proportion changes in MDD and remission and suggest instead distinct cellular functional changes in MDD: Progression over MDD episodes may specifically disrupt cortical processing and associative functions mediated by superficial layer IT neurons, while MDD state changes broadly affect both output (ET) and internal (IT) cortical circuits, inhibitory interneurons, and endothelial cells linked to ECM dysregulation.

### Up- and downregulated genes distinguish MDD phases and broader disease links

To determine whether the DEGs involved in progressive and state contrasts are implicated in symptom dimensions and other disorders beyond depression, we analyzed their enrichment (p-value < 0.01) in disease-specific gene sets independently identified in the DisGeNET [17] and PsyGeNET databases [18]. Figure 3 illustrates the association between the upregulated (yellow) and downregulated (blue) DEGs from all contrasts with gene sets associated with various diseases or symptoms, grouped by selected, clinically relevant semantic category on the y-axis.

**Figure 3:**
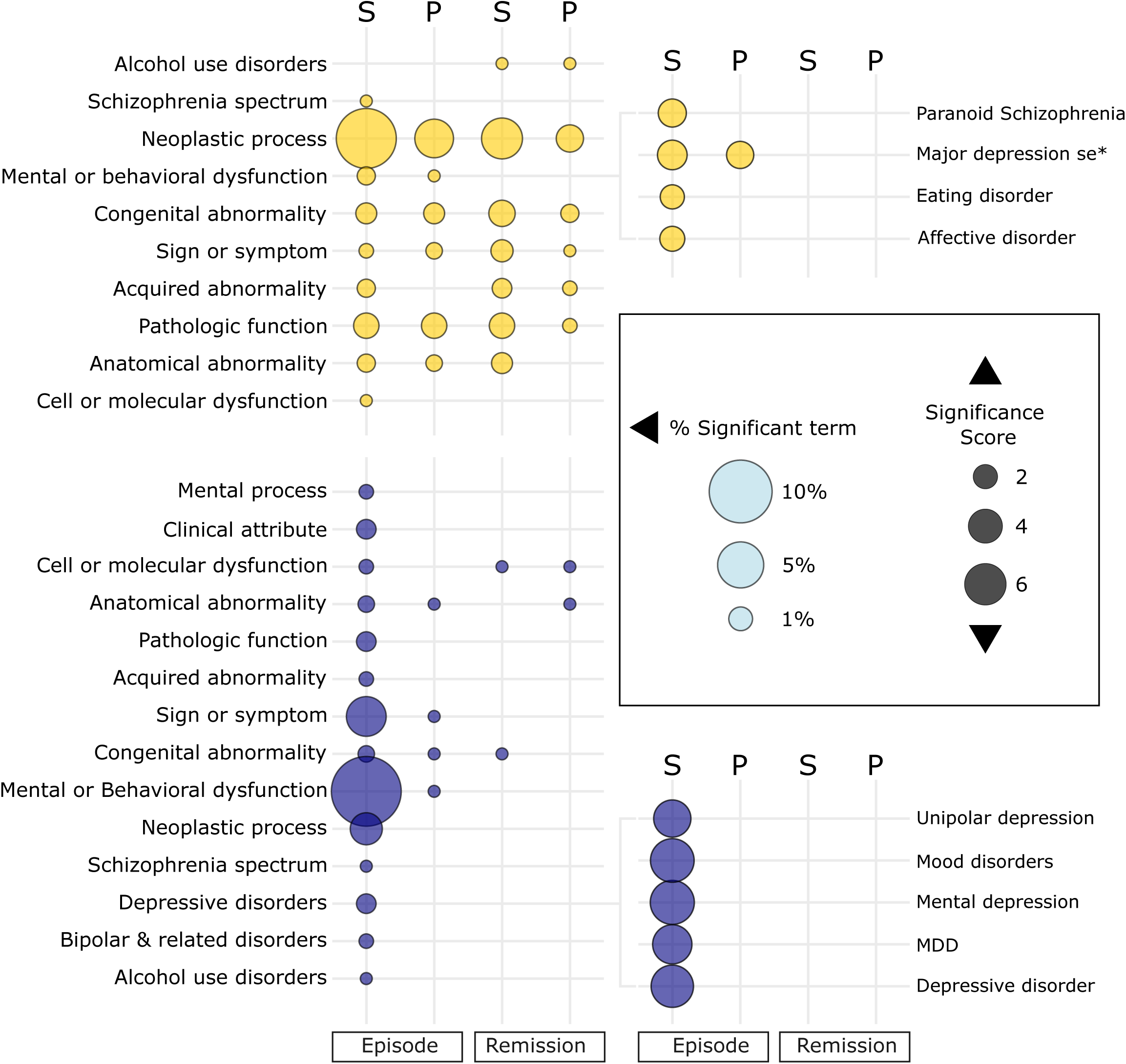
Disease associations of DEGs across MDD phases: Enrichment of upregulated (yellow, top) and downregulated (blue, bottom) DEGs in disease-associated gene sets from DisGeNET and PsyGeNET. The left-side labels represent disease semantic categories. In the main plot (left), circle size is proportional to the percentage of significant disease terms within each disease semantic category, calculated as: (number of contrast specific significant disease terms within a given semantic category / total number of significant disease terms across all contrast [=807]) × 100. In the insets (right), circle size is proportional to the significance score, calculated as -log10(p-value). Labels for the insets indicate the specific diseases corresponding to the disease semantic categories shown in the main plot. Note that for most disease semantic categories, the number of diseases per contrast is less than 1%. See Supplementary Table S7 for full details on the disease-specific enrichment analysis and disease semantic categories.

Among the downregulated DEGs (Figure 3, bottom), those from state-episode were associated with disease semantic terms related to psychiatric illnesses (*schizophrenia spectrum, depressive disorders, bipolar & related disorder*) and comorbid conditions (*alcohol use disorder (AUD) and mental processes*), with the strongest association found in *mental and behavioral dysfunction*. Notably, these DEGs were exclusively linked to *depressive disorders*, suggesting that downregulated DEGs from episode-state-changes may serve as a signature for MDD. Interestingly, the downregulated DEGs from progressive episodes, aside from a few diseases linked with *mental and behavioral dysfunction* (*autism spectrum disorder and mild cognitive disorder*), showed no significant associations with psychiatric or related disorders, suggesting that progressive changes contribute differently from state changes to MDD pathology. In contrast, downregulated DEGs from both state and progressive remission showed minimal association with disease semantic terms, suggesting a relative molecular decoupling from molecular mechanisms associated with disease during remission.

In contrast, the upregulated DEGs (Figure 3, top) were more evenly associated with multiple semantic terms across both state and progressive contrasts during episode and remission. Given that episode and remission represent opposite biological phenomena, this suggests that the neurobiological adjustments that occur during remission also occur during episodes, potentially as a compensatory mechanism aimed at restoring normal function. Additional changes were specific to either the episodic or remission phases. For example, episodic DEGs from state and progressive contrasts were uniquely linked to *mental or behavioral dysfunction* with *major depression* (*single episode*) being common to both. Similarly, DEGs from state and progressive remission contrasts were exclusively linked to *alcohol use disorder*, consistent with reports of mood disorder and substance use disorder comorbidity during remission [31–33]. Notably, DEGs from all four contrasts showed a strong association with the semantic term related to *neoplastic process* and *pathologic function*, involving disorders that influence the ECM and angiogenesis— dysregulations that were also highlighted in our pathway analysis for both state-episode and state-remission.

Taken together, these findings suggest that downregulated genes from state-episode changes may serve as a signature for MDD, while upregulated genes across both contrasts may reflect compensatory mechanisms during both episodes and remission.

### Genetic association distinguishes trait- and state-dependent MDD symptoms

To investigate the genetic relevance of transcriptomic changes across the MDD trajectory, we performed gene-set enrichment analysis using MAGMA [23] on summary statistics from genome-wide association studies of MDD [25] and those focusing on MDD ascertainment and symptom structures [26]. We assessed the association of DEGs from each contrast using two complementary approaches: a self-contained approach (Figure 4, colored bars, q-value < 0.05), which evaluates each DEG set independently for its overall association with MDD, and a competitive approach (Figure 4, black lines, p-value < 0.05), which compares the association of each DEG set to all other genes across the genome to determine its distinct relevance to MDD.

**Figure 4:**
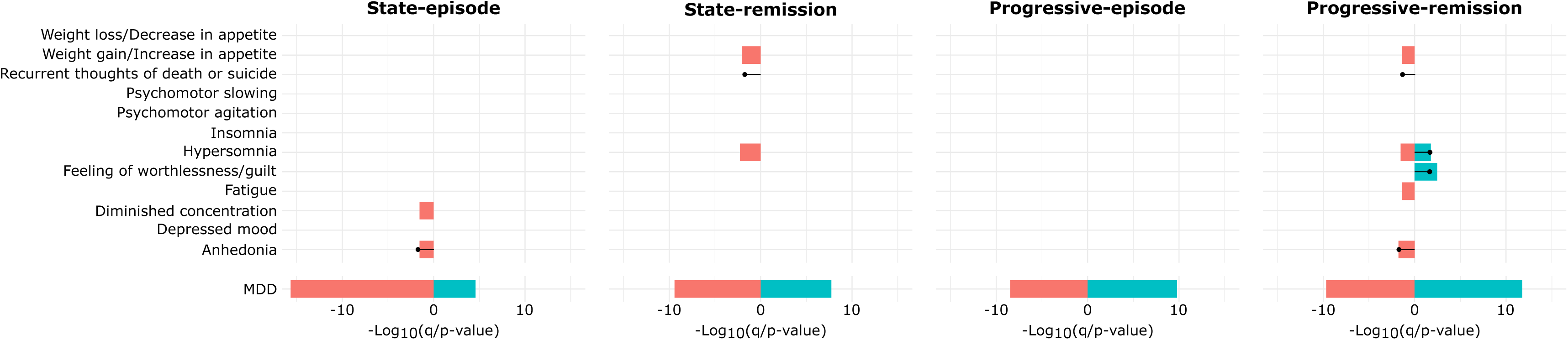
Genetic association of DEGs with MDD and symptom manifestations: Gene-set enrichment analysis using MAGMA linking DEGs from state and progressive contrasts to MDD genome-wide association study summary statistics. The colored bars represent the self-contained approach, which tests each DEG set independently, while the black line represents the competitive approach, which evaluates DEG associations relative to all genes in the genome. The plots for self-contained and competitive approaches were generated using the same x-axis scales and overlayed for better comparison.

In the self-contained analysis, all eight DEG sets (spanning state vs. progressive contrast, episode vs. remission phase, and up-vs. downregulation) showed highly significant associations with MDD (Figure 4, bottom rows), indicating that transcriptomic changes across the disease trajectory are highly consistent with genetic findings in MDD. Within the MDD ascertainment and symptom structures, state-episodes were associated with terms of diminished concentration and anhedonia, whereas progressive-episodes showed no such associations. In contrast, remission phases were linked to weight gain/increased appetite and hypersomnia in both state and progressive contrast. Additionally, progressive remission was uniquely associated with terms of feelings of worthlessness/guilt, fatigue, and anhedonia.

In the competitive analysis, remission in both state and progressive contrasts were associated with recurrent thoughts of death and suicide. Consistent with the self-contained analysis, progressive remission was also linked to hypersomnia and feelings of worthlessness/guilt. Notably, anhedonia was associated with both state-episode and progressive-remission.

Overall, this analysis suggests that our transcriptomic findings are genetically linked to MDD and its symptom manifestations, including those observed during remission. In particular, genes associated with anhedonia may reflect a persistent disease trait, present across both episodes and remission.

### Molecular mechanisms of drug actions are highly represented in DEGs associated with MDD pathology and recovery

The signature-matching principle in drug repurposing posits that drugs mimicking disease-specific gene expression profiles may act as pro-disease agents, whereas those reversing such profiles may exert therapeutic effects [34]. Based on our findings that downregulated genes mark disease states and upregulated genes mark compensatory changes (Figure 3); we compared each contrast-specific DEG set with drug-induced upregulated expression profiles. Drugs whose signatures opposed the MDD downregulated genes were identified as potential therapeutics, while those aligning with the MDD upregulated genes were considered pro-disease (q-value < 0.05). Several drugs with known therapeutic (Creatine, D-Serine, Morphine, Fluoxetine, Levetiracetam, pioglitazone and clozapine) [35–53] or pro-disease (Cadmium, Soman, Lipopolysaccharide, Dexamethasone, Interferon Beta-1A and Isotretinoin) [54–69] effects, confirmed through literature review, emerged from the analysis (Figure 5A). Among these, fluoxetine (antidepressant) [70], dexamethasone (glucocorticoid) [71] and lipopolysaccharide (inflammatory agent) [59] exemplify known therapeutic and pro-disease mechanisms, supporting the validity of our approach.

**Figure 5:**
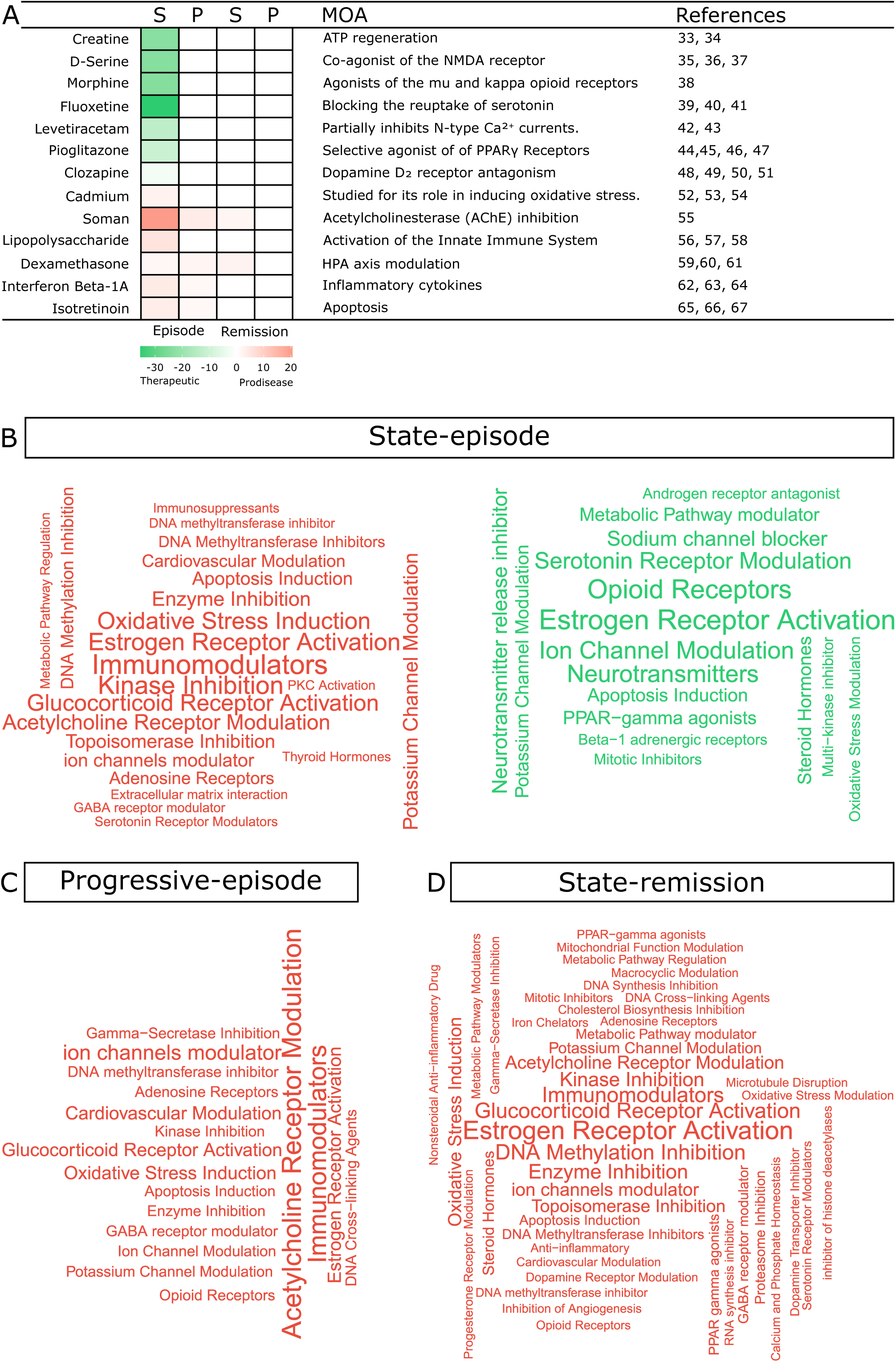
Molecular mechanisms linking therapeutic and pro-disease effects in MDD: **(A)** Enrichment analysis of contrast-specific DEGs in drug-induced upregulated expression profiles. Enrichment of downregulated DEGs indicates potential therapeutic effects (green), whereas enrichment of upregulated DEGs suggests pro-disease effects (red), based on the signature-matching principle. Selected drugs with known mechanisms of action (MOA) are annotated with relevant references. **(B-D)** A word cloud displays the MOA for pro-disease (red) and therapeutic (green) drugs across contrasts. The size of each term corresponds to its frequency within the respective contrast. See Supplementary table S8 for full drug-matching results.

To summarize the mechanisms of action (MOA) of the identified drugs, we show a word cloud-based semantic enrichment for pro-disease (red) and therapeutic (green) drugs across contrasts. For state-episode, a complex network of pro-disease mechanisms (Figure 5B, red) emerged, encompassing neurotransmitter and receptor modulation (*acetylcholine, adenosine, GABA, serotonin, estrogen, glucocorticoid and thyroid hormones*), ion channel regulation (*including potassium channel modulation*), immune dysregulation (*immunomodulators and immunosuppressants*), epigenetic disruptions (*DNA methylation inhibition and topoisomerase inhibition*), metabolic and cardiovascular modulation, extracellular matrix interaction, apoptosis induction and oxidative stress, together consistent with the multi systems pathologies of MDD. In contrast, therapeutic mechanisms (Figure 5B, green) were primarily centered on modulating hormone signaling (*androgen receptor antagonism, estrogen receptor activation, steroid hormones, and PPAR-gamma agonists*), regulating neural excitability and neurotransmitter function (*beta-1 adrenergic receptor modulation, ion channel modulation, sodium channel blockade, opioid and serotonin receptor modulation, neurotransmitter release inhibition*), controlling cell proliferation and survival (*mitotic inhibitors, multi-kinase inhibitors, apoptosis induction, oxidative stress modulation*), and adjusting metabolic pathways. Notably, apoptosis induction, estrogen receptor activation, oxidative stress modulation, and potassium channel modulation appeared in both pro-disease and therapeutic categories, suggesting that their effects may depend on dosage, cell-type or disease state, reflecting a pleiotropic nature within MDD pathology.

For progressive episodes (Figure 5C, red), several of the pro-disease mechanisms—glucocorticoid receptor activation, oxidative stress, immune modulation, ion channel dysregulation, and epigenetic alterations—seen in episode state persisted, indicating that these pathways may play a foundational role in disease progression, contributing to both the onset and long-term development of MDD. Additionally, mechanisms such as DNA cross-linking, gamma-secretase inhibition, and kinase inhibition emerged specifically in progressive episodes, potentially reflecting molecular disruptions that evolve as the disorder advances.

Consistent with their decoupling from the disease state, no significant therapeutic targets were identified during state-remission. However, several pro-disease targets (*apoptosis induction, estrogen receptor activation, metabolic pathway regulation, potassium channel modulation and serotonin receptor modulation*; Figure 5D, red), which in this context may represent pro-remission mechanisms, overlapped with both pro-disease and therapeutic mechanisms observed in the state contrast (Figure 5B, red and green), suggesting that some pathways promote remission through homeostatic processes, while others may drive treatment-resistant depression or relapse.

Overall, these findings suggest that there are narrow therapeutic targets and widespread pro-disease mechanisms. They also highlight the dynamic nature of molecular pathways in MDD, where the same mechanisms can either exacerbate or alleviate disease depending on dose, disease state, or timing.

## Discussion

In this study, we systematically examined transcriptomic signatures across distinct clinical phases of MDD—episodes and remission—to distinguish progressive from state-specific molecular changes. Building upon our previous work [4] that identified oscillatory transcriptomic patterns across MDD states, we now extend this framework to dissect how progression during episode or remission compares with state-specific changes in gene expression, cellular vulnerability, disease associations, genetic risk, and pharmacological targeting.

DEGs and affected biological pathway enrichments across state and progressive contrasts (Figure 1C-E) enabled distinction between biological processes associated with acute disease burden from those that accumulate gradually with illness progression. ECM-related pathways were consistently upregulated in both episode and remission, whereas catalytic and organelle-related pathways were selectively restored during remission. These findings suggest a dual nature of MDD remission: partial molecular recovery alongside persistent ECM disruption, implicating ECM dysregulation as both trait-like feature and progressive molecular hallmark of the disorder. Notably, persistent ECM-pathway upregulation across episode and remission aligns with evidence from MDD models [72, 73], where ECM remodeling—particularly involving perineuronal nets—contributes to stress-induced synaptic rigidity and impaired plasticity. Their continued activation during remission suggests a potential role in treatment resistance and relapse, warranting further investigation.

Selective transcriptional alterations in sgACC cell types (Figure 2) reveal that specific neuronal populations exhibit heightened vulnerability in MDD, despite stable overall cell-type proportions. Progressive changes were primarily confined to superficial-layer IT pyramidal neurons, while state-dependent alterations extended to deeper-layer pyramidal neurons and SST-expressing interneurons. This supports the notion that associative cortical circuits are gradually impaired with disease progression, and broader cortical networks are impacted by state-specific disease burden. These observations are consistent with prior work demonstrating that layer 2/3 pyramidal neurons are critical for stress modulation—such as the disruption of *Wfs1* in these neurons leading to exaggerated stress responses and HPA axis hyperactivity [74]. The involvement of SST interneurons also aligns with converging human [4, 27, 28] and animal [75] studies indicating that SST-positive GABAergic deficits disrupt cortical processing and contribute to depressive phenotypes.

In parallel, upregulated genes during both episodes and remission were enriched in endothelial cells, suggesting neurovascular dysfunction as underappreciated contributor to MDD. Notably, remission states did not exhibit neuron-specific changes but showed enrichment of upregulated genes in endothelial cells, implicating neurovascular processes in recovery. Because endothelial cells form the front line of blood-brain barrier (BBB), their activation aligns with evidence showing chronic stress-induced BBB disruption via VEGF signaling [76, 77].

Importantly, many of these endothelial-associated changes converge on ECM pathways, which remained persistently upregulated in both episode and remission phases (Figure 1D & E). Given the ECM’s critical role in maintaining BBB integrity [78], synaptic stability [79], and perineuronal net formation [80], its persistent dysregulation may represent a shared molecular axis linking neuronal vulnerability, vascular dysfunction, and incomplete remission. These observations suggest a model in which MDD progression is driven by sequential neuronal circuit impairment, while remission involves partial functional restoration constrained by enduring neurovascular-ECM pathology—potentially contributing to treatment resistance and relapse.

Complementing the cell-type-specific findings, transcriptomic alterations across MDD phases also reflected a strong genetic basis. DEGs associated with the state and progressive contrasts were significantly enriched for genetic loci implicated in GWAS of MDD and related traits. Of particular interest, anhedonia, a hallmark symptom of MDD [81], was associated with both state-episodes and progressive-remission DEGs, positioning it as a persistent, trait-like component of the disorder. This is consistent with studies highlighting shared genetic architecture between anhedonia and multiple psychiatric disorders, including schizophrenia and bipolar disorder [82]. Interestingly, our transcriptomic associations linked symptoms typically associated with atypical depression—such as hypersomnia, increased appetite, and feelings of guilt [83]—to the remission phases. Atypical depression is characterized by mood reactivity, where symptoms temporarily improve in response to positive events [83–85], overlapping conceptually with the partial symptom resolution observed during remission. While this connection warrants further investigation in longitudinal clinical studies, the presence of these symptoms during remission in our dataset may reflect residual pathology or serve as prodromal indicators of relapse.

Differential associations of DEGs with disease-specific gene sets and drug-induced expression profiles underscore the complementary roles of down- and upregulated genes in MDD pathology. While transcriptomic analyses capture the directionality of gene expression changes, they do not inherently designate which alterations mark disease state versus compensatory adaptation. Our findings suggest that downregulated genes, particularly in the state-episode, are more specifically associated with depressive disorders and may serve as molecular signatures of disease burden. In contrast, upregulated genes were broadly enriched across both episode and remission states and spanned multiple disease categories, consistent with a putative compensatory or adaptive role. This interpretation was further validated through a signature-matching drug repurposing approach, which revealed that fluoxetine—an established antidepressant [70]—opposed the downregulated gene signatures, while dexamethasone and LPS—agents associated with stress [71] and inflammation-induced depression [59]—mirrored the upregulated patterns. Mechanistically, both therapeutic and pro-disease compounds act through overlapping biological pathways, including oxidative stress modulation, estrogen signaling, and ion channel activity. These findings align with prior evidence demonstrating that pleiotropic and context-dependent nature of molecular pathways in MDD [86], wherein the same mechanism can support either recovery or pathology depending on cell type, dosage or disease phase. While drug repurposing based on transcriptional signature can serve as a tool for elucidating disease mechanisms, we caution that, for uncovering candidate therapeutics, interpretation of results may require contextualization through disease-linked gene associations to discern pathogenic versus adaptive molecular changes.

This study has inherent limitations associated with human postmortem research. First, while our uniquely staged cohort enabled the investigation of MDD progression from episode to remission, the rarity of such samples limited our ability to perform external validation in an independent dataset, particularly for progressive contrasts. Nevertheless, our findings showed strong internal consistency, robust alignment with prior studies, and significant overlap with GWAS summary statistics, providing genetic-level validation of the disease relevance of our transcriptomic signals. In addition, we confirmed key gene expression patterns using qPCR, demonstrating technical concordance with the RNA-seq measurements. Second, while our approach captures molecular progression by comparing individuals across disease stages, true longitudinal tracking within subjects is not feasible in postmortem cohorts. Therefore, the progressive changes described here should be viewed as hypothesis-generating, warranting future validation in longitudinal or functional models. Third, within the limitations of postmortem studies, some comparisons were inherently imbalanced in sample size due to sample availability, and the overall sample size was underpowered for rigorous sex-specific analysis.

Future work with larger and more balanced cohorts will be needed to address these limitations and to explore potential sex differences.

## Acknowledgments

This work is supported by Institutional Development Award from the National Institute of General Medical Sciences of the National Institutes of Health under Grant # 2P20GM103432 and 2P20GM121310 to RS.

