## Supplementary material for "Molecular Characterization of the Progressive Landscape of Depression": Sharma_Supplementary_Information_and_Notes_Revision-02.docx

**This PDF file includes:**

Supplementary Figures S1 to S4 and related notes

Supplementary Table S1 to S8 notes

References

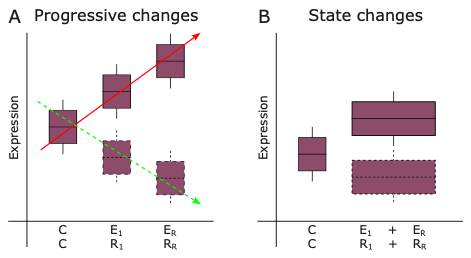

**Supplementary Figure S1: Progressive and state contrasts to characterize MDD transcriptional dynamics.** Progressive contrast (A) models a cross-sectional linear trend of upregulated (red) or downregulated (green) gene expression across clinical stages using ordinal groups for both episodes [Control (C) → First episode (E_1_) → Recurrent episode (E_R_)] and remissions [Control (C) → First remission (R_1_) → Recurrent remission (R_R_)]. This approach aims to identify genes whose expression changes gradually and consistently with illness trajectory at the group level, capturing the slope of change and revealing dose-response–like patterns along disease progression rather than within-subject longitudinal changes. In contrast, state contrast (B) compares all affected individuals (either in episode or remission) directly to controls, identifying genes consistently dysregulated regardless of disease progression. This contrast captures average differences and highlights state-dependent dysregulation independent of progression. Upregulated genes are shown with solid boxes, and downregulated genes with dotted boxes.

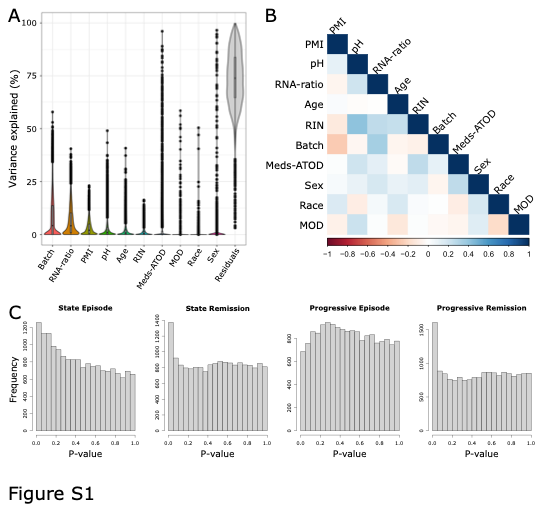

**Supplementary Figure S2: Covariate assessment and quality control for differential expression analysis.** **(A) Violin plot showing the distribution of variance in gene expression explained by each variable after filtering out low-expressed genes. Each violin represents a different variable, with the y-axis depicting the proportion of variance in gene expression attributable to that variable. (B)** Covariate correlation heatmap assessing multicollinearity among batch effects, RNA integrity, postmortem interval (PMI), brain pH, age, sex, race, and medications at the time of death. No highly correlated variables were observed, ensuring that covariate correction did not over-adjust biologically meaningful variance. **(C)** P-value histograms for differential expression analysis, demonstrating uniform distribution, indicating appropriate statistical control.

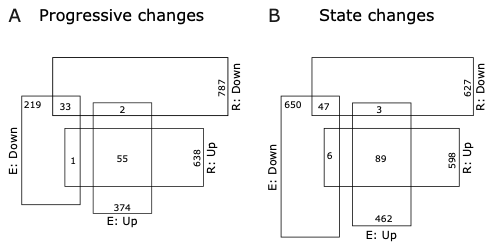
**Supplementary Figure S3: Minimal overlap between episode- and remission-specific DEGs across contrasts.** Venn diagrams illustrate minimal overlap of DEGs between episode and remission states within both progressive **(A)** and state **(B)** contrasts, based on a significance cutoff of p-value <0.05. This indicates that episode and remission represent separate phases of illness.

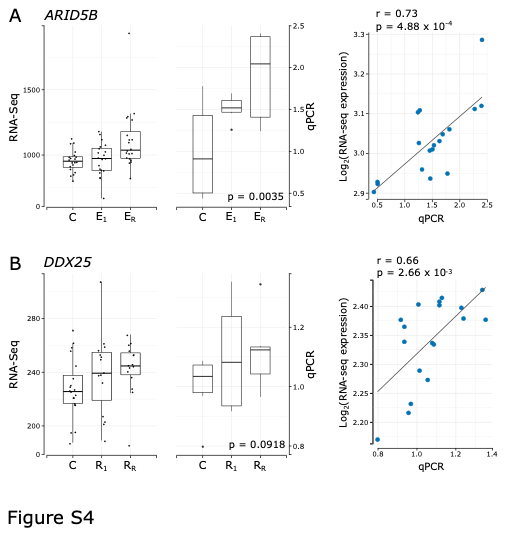

**Supplementary Figure S4: Technical validation of RNAseq findings using qPCR.** To demonstrate that RNA-seq expression patterns could be reproduced by an independent assay, we selected (A) *ARID5B* and (B) *DDX25* as representative genes from the progressive-episode and progressive-remission contrasts, respectively. Both genes showed nominally significant changes within their respective contrasts and were key contributors to the associated GO-term enrichment signals (i.e., part of the gene overlap accounting for the enrichment signal). Specifically, *ARID5B* was associated with the GO term “cell migration,” mapping to the broader theme of “cell motility,” while *DDX25* was associated with the GO term “catalytic activity, acting on a nucleic acid,” mapping to the broader theme of “catalytic activity.” qPCR performed on the same samples showed strong concordance with the RNA-seq measurements, supporting the technical reliability of the RNA-seq data.

Notably, ***ARID5B* encodes a chromatin-associated transcriptional regulator that modulates enhancer accessibility and inflammatory or metabolic programs relevant to stress-related disorders [1]. Additionally, variants in this gene were recently identified as candidate causal contributors to MDD in a large-scale exome sequencing study [2].** In this context, altered *ARID5B* expression could affect stress- and cytokine-responsive transcriptional states in cortical regions, providing a plausible link to episode-related molecular burden.

**By contrast, *DDX25* has not been previously associated with depression, which aligns with our observation that its signal was specific to remission rather than active episodes.** However, given its known role in RNA processing [3, 4], the remission-associated *DDX25* expression pattern observed in our study may reflect compensatory adjustments in RNA metabolism accompanying recovery.

**Supplementary Table S1: Demographic information of postmortem sample:** Postmortem brain samples were collected from control subjects (**Group I**, n = 20) and four major MDD cohorts: **single episode** (Group II*,* n = 20), **single episode remission** (Group III*,* n *=* 15), **recurrent MDD** (Group IV*,* n = 20), and **recurrent remission** (Group V*,* n = 15). Each subject’s **DSM-IV diagnoses, mode of death (MOD), cause of death (COD), and demographic features** (sex, age, race, postmortem interval [PMI], brain pH, RNA integrity metrics) are detailed. MDD groups were classified based on **disease severity, recurrence, and remission status**. The dataset includes information on **medications at time of death (Meds ATOD), tobacco and substance use, depression history (Dep ATOD), duration of illness (DOI), recurrence, history of psychosis, and remission status.** Control subjects with a DSM-IV diagnosis are included, and their **DOI is provided in parentheses.**

At the bottom, **summary statistics** report the **mean and standard deviation for key biological variables** (age, PMI, pH, RNA ratio, RIN) for each group. **Pairwise comparisons** (p-values) assess differences between groups across these biological variables, confirming that there were **no significant differences** (p-value > 0.05 for most comparisons), ensuring that these factors do not confound subsequent transcriptomic analyses.

**Supplementary Table S2: Cohort Statistics:** Summary of demographic and biological characteristics of the study cohorts. The table presents mean values for age, postmortem interval (PMI), brain pH, RNA ratio, and RNA integrity number (RIN), along with percentages for sex, suicide cases, and antidepressant use at the time of death (ATOD). Notably:

1. PMI, pH, RNA ratio, and RIN values are comparable across groups, minimizing confounding effects.
2. Suicide rates were observed only in MDD groups, with 50% in single-episode MDD and 40% in recurrent MDD, while remission groups showed no suicide cases.

**Supplementary Table S3:** DEGs identified across state and progressive contrasts during both episode and remission phases. The table includes gene-level statistics from DESeq2 analysis, with columns representing:

1. **Gene stable ID:** Unique identifier for each gene
2. **Gene description:** Functional annotation of the gene
3. **Gene name:** Common gene symbol
4. **baseMean:** Average expression level across all samples
5. **log2FoldChange:** Magnitude and direction of differential expression
6. **lfcSE:** Standard error of the log2 fold change
7. **stat:** Wald statistics for differential expression
8. **p-value:** Statistical significance before multiple testing correction
9. **padj:** Adjusted p-value (FDR-corrected) for significance threshold

DEGs were identified separately for state episode, state remission, progressive episode, and progressive remission contrasts. Based on nominal significance (p-value < 0.05), 7.59% of tested genes were significant in the state episode contrast, 7.97% in the state remission contrast, 4.13% in the progressive episode contrast, and 8.82% in the progressive remission contrast.

**Supplementary Table S4:** Pathway enrichment analysis for DEGs across state and progressive contrasts during both episode and remission phases. This table represents all pathways associated with different DEG contrasts, while Figure 1C-E highlights selected ontological groups from this dataset.

- **Pathway Significance Scores:** Columns for **State Episode, State Remission, Progressive Episode, and Progressive Remission** display significance scores calculated as **-log_10_(q-value)** for all significant pathways (**column B**). **Negative values** indicate **downregulation**, while **positive values** indicate **upregulation**.
- **Ontological Groupings**: Boolean values (TRUE/FALSE) indicate whether a given pathway falls under broader functional categories, including epithelium, angiogenesis, synapse, axon, immune system process, glia, cell motility, catalytic activity, organelle, and extracellular matrix (ECM). These classifications correspond to the color-coded themes in Figure 1.

**Supplementary Table S5 and S6:** Deconvolution analysis using Support Vector Regression (SVR) [5] and Ordinary Least Squares (OLS) [6] methods were performed to estimate cellular composition across study conditions in episode (Table S5) and remission (Table S6) states. The tables display the estimated proportions for each sample, representing the fraction of the bulk RNA-seq signal attributable to each reference cell type. Columns g0–g24 correspond to the different reference-derived cell-type signatures, and rows represent individual samples from Control, Single Episode/Remission, and Recurrent Episode/Remission groups.
The negative values in the tables reflect model-fitting artifacts that are standard in reference-based deconvolution analyses and do not represent biologically meaningful negative proportions. The last two rows, highlighted in orange, indicate that all results were non-significant (NS) for state and progressive contrasts, confirming that no significant differences in cellular composition were observed between groups. Human ACC single-nucleus RNA-seq data were obtained from the Allen Brain Atlas [7]. Cell-type clustering and summarized cell-specific expression data were obtained using the Seurat pipeline.

**Supplementary Table S7:** Disease enrichment analysis for DEGs across state and progressive contrasts during both episode and remission phases. This table represents all diseases associated with different DEG contrasts, while Figure 3 highlights selected disease categories from this dataset.

- **Disease Significance Scores:** Columns for State Episode, State Remission, Progressive Episode, and Progressive Remission show significance scores computed as -log_10_(q-value), where negative values indicate enrichment of downregulated DEGs, and positive values indicate enrichment of upregulated DEGs.
- **Disease Classification:** The PsyGeNet and DisGeNet columns indicate whether a disease is reported in these databases. The Disease Semantics column categorizes diseases into broad classifications, such as neoplastic processes, congenital abnormalities, and mental and behavioral disorders, among others.
- **Pathological and Clinical Associations:** Boolean values (TRUE/FALSE) indicate whether a disease is linked to specific pathological or clinical attributes, including psychiatric disorders (e.g., alcohol use disorder, schizophrenia, depressive disorders), congenital abnormalities, neoplastic processes, and laboratory findings. These attributes correspond to the y-axis in Figure 3.

**Supplementary Table S8:** Drug-specific analysis was performed to assess the enrichment of DEGs derived from state and progressive contrasts in both episode and remission states within the upregulated gene signatures of drug responses from human or mouse cell lines. Selected drugs with known therapeutic and pro-disease effects are highlighted in figure 5A, as an internal validation of the approach.

**Drug Significance Scores:** Columns for State Remission, Progressive Remission, State Episode, and Progressive Episode represent significance scores computed as -log_10_(q-value). Negative scores indicate that downregulated DEGs are enriched in the upregulated drug response signatures, suggesting a therapeutic effect according to the signature-matching principle. Likewise, positive scores indicate that upregulated DEGs are enriched in these signatures, suggesting a pro-disease effect.

**Therapeutic vs. Pro-Disease Effects**: Boolean values (TRUE/FALSE) indicate whether a drug exhibits a therapeutic or pro-disease effect in each condition, based on the signature-matching principle, and are categorized under State and Progressive contrasts for both Episode and Remission states.

**Drug Information**: The Drug column lists the tested compounds, with Classification indicating their broader pharmacological category (e.g., neurological drugs, anti-inflammatory agents), and MOA (Mechanism of Action) describing their molecular targets, such as opioid receptors, immune system modulators, or estrogen receptors. MOAs were identified through literature search and public database annotations or inferred from the rationale described in the respective GEO study. For example, heavy metals tested for their role in inducing oxidative stress were accordingly assigned “oxidative stress” as their MOA. The MOA column was used to generate the word cloud in Figure 5B.
